# Protein quality of *Hyptis spicigera* syn. *Cantinoa americana* (*amola*): a locally valued yet underutilized African food crop adaptable to an uncertain future

**DOI:** 10.1101/2025.06.20.660461

**Authors:** Eliot T. Masters

**Affiliations:** Independent researcher

**Keywords:** Plant-based nutrition, on-farm biodiversity, traditional food plants, neglected food plants, orphan crops, plant protein, precolonial agriculture, plant domestication

## Abstract

Food security and nutrition are increasingly slipping the grasp of those most vulnerable to market shocks, regional conflict and the effects of climate change. For smallholder farmers of sub-Saharan Africa, crop biodiversity can mitigate nutritional and livelihood risks, provide a varied diet and reinforce systemic resilience through redundancy of functions. However, ongoing and accelerating processes of cultural and genetic erosion have profoundly diminished varietal diversity, risking the loss of locally valued food plants from future harvests. Presented here is new data on the compositional profile of *Hyptis spicigera* Lam. syn. *Cantinoa americana* (Aubl.) Harley & J.F.B.Pastore (Lamiaceae), an underutilized and profoundly neglected precolonial food crop domesticated in historical times and once widely cultivated across equatorial Africa for its protein-rich oilseed. The amino acids profile of hyptis seed has not previously been determined, nor has its protein quality been assessed.

**Objectives:** The purpose of the study was to investigate the nutritional value of hyptis seed including its proximate composition, amino acids profile and protein quality. A secondary objective was to construct a comprehensive review of historical literature and secondary data on the species, for which the published record is sparse and antiquated.

**Methods:** A blended sample of hyptis seed was obtained from Apur market at Adwari subcounty in the Otuke District of northern Uganda. Proximate nutritional composition was assessed in three replicates, and amino acid analysis was performed in four replicates for 19 amino acids. A protein quality test was conducted using the WHO/FAO Protein Digestibility-Corrected Amino Acid Score (PDCAAS) method. All relevant historical literature was consulted, and a comparative review undertaken of flower morphology in 138 digitized herbarium specimens.

**Findings:** Proximate composition of the seed was found to comprise 28.8% lipids and 19.7% protein, a lipid to protein ratio of 3:2 comparable to some *Sesamum* species. The nutritionally limiting amino acid was found to be lysine, although the deficiency is not statistically significant based on an amino acid score of 99% and a standard error (SD) of 0.18. Protein quality was found to be relatively high, with a PDCAAS of 84% comparing favorably both to pulses (pea) and pseudocereals (quinoa). An historical review of the species valorizes hyptis as a valued and productive resource within smallholder farming systems, a precolonial African food crop meriting further attention.

## 1.0 Introduction

Food security and nutrition are increasingly slipping the grasp of those most vulnerable to market shocks, regional conflict and the effects of climate change. Currently 280 million people in 59 countries face acute food and nutritional insecurity (FSIN and GNAFC, 2024). In small-scale farming systems, rising temperatures and irregular rainfall are driving down crop yields, with further losses anticipated without effective regional adaptation (Lobell & Field, 2007; Cohn et al., 2017; Zhao et al., 2017; Williams et al., 2018; Zhu et al., 2022). Future productivity will be further compromised by the effects of increasing temperatures on soil biota (Soares et al., 2019), while increasing atmospheric carbon inhibits crop nitrogen uptake, thus reducing the protein content and nutritional quality of cultivated foods (Taub et al., 2008; Lumsden et al., 2024).

Agricultural progress has relied upon the timely adoption of plant species and varieties which respond effectively to a changing climate (Vaughan et al., 2007), and increasingly rapid breeding and varietal replacement are seen as strategically critical within smallholder cultivation (Atlin et al., 2017). However, for varietal adaptation to remain viable will depend upon the continued existence of a viable resource base of crop varieties, maintained in cultivation and exchanged through farmer-to-farmer interaction (Mijatović et al., 2013; Zabel et al., 2021). A growing consensus underlines the urgency of preserving traditional food crops and varieties which are locally or regionally significant but lacking in wider recognition (variously known as underutilized, neglected or ‘orphan’ crops), against an accelerating trend of cultural and genetic erosion whereby technical knowledge and germplasm are lost (Talabi et al., 2022; Dempewolf et al., 2023 ; McCouch & Rieseberg, 2023).

At the same time, baseline data on the nutritional value of these diminishing resources is often lacking (Padulosi et al., 2013; Baldermann et al., 2016). Dietary protein in particular has been identified as a particularly important macronutrient given its central role in regulation of cellular metabolism, growth and development and (Schönfeldt & Hall, 2012; Nolden & Forde, 2023). Global demand for edible protein has been increasing in recent decades, with increasing interest in plant-based proteins, protein diversification and the emergent concept of protein security (Langyan et al., 2022; Rzegotta, 2024; Kakabouki et al., 2025).

However, human nutritional requirements derive from protein quality, a measure of nutrient density based on a complete profile of the essential amino acids (Beal et al., 2024). Since plant proteins tend to be limiting in one or more of these - cereals in isoleucine, lysine, threonine, and tryptophan; legumes in methionine - complementary protein sources must be combined in order to avoid protein deficiency (Schönfeldt & Hall, 2012; Langyan et al., 2022). Recent studies stress the need for assessment of protein quality as a critical determinant of nutrient adequacy of foods and diets (Pikosky et al., 2022; Nolden & Forde, 2023; Beal et al., 2024), but most recent studies on plant protein quality have been limited to higher-profile leguminous crops (*e.g.* Tanimonure et al., 2023; Singh et al., 2023) or pseudocereals such as amaranth (Mota et al., 2016; Vidaurre-Ruiz et al., 2023), while some underutilized crops of local value have remained to be investigated.

*Hyptis spicigera* Lam. (syn. *Cantinoa americana* (Aubl.) Harley & J.F.B.Pastore) Lamiaceae is one such traditional crop of cultural significance in northern Uganda, where it is esteemed as the traditional delicacy *amola*.

In Uganda, hyptis was widely cultivated as a major food crop in the Acholi lands on the western banks of the Nile, where it was first observed during the early dry season of November 1862 as one of the three grain crops identified by Grant (1864) alongside finger-millet (*Eleusine coracana* (L.) Gaertn.), and a *Hibiscus* species (*H. sabdariffa* L.) – the source of another highly specialized and labor-intensive seed protein crop, known locally as *toke* (Epila-Otara, 2013). These early reports were corroborated during a 1905 collecting mission for the Royal Botanic Gardens at Kew by the botanist Morley Dawe, who reported food prepared from hyptis to be “much relished” (Stapf et al., 1906). Hyptis was listed by Driberg (1923) and Tarantino (1943, 1949) among the earliest food crops cultivated by the Lango. Purseglove (1943) found the crop under cultivation in the West Nile District, “for its seeds, which are rich in oil, used in the same way as [sesame], and often eaten with green vegetables.”

Celebrated in poetry and prose by the renowned Acholi poet Okot p’Bitek as a deep cultural signifier of the ancestral village (1983, 1986), hyptis was identified as a minor tropical food crop by Purseglove (1988), and was listed among the cultivated edible plants of by Goode (1989) and by Katende et al. (1999). More recently, hyptis has been identified as a notable Ugandan plant genetic resource for food and agriculture (NARO, 2008), was found to be traded in the markets of western Bunyoro region (Agea et al., 2013), and was recognized as a currently significant food to the Acholi (Nyero et al., 2021) and the Lango (Masters, 2021).

As described in some detail by the author in that prior publication, the dish called *amola* is customarily prepared for special guests and on festive occasions, reflecting its limited production and the arduous tasks involved in its preparation (Epila-Otara, 2013). The hyptis seed (more accurately, a botanical fruit) is small and hard, and when can be conserved for several years in storage (Masters, 2021). Preparing *amola* has traditionally involved stone-grinding the lightly roasted seed into a flour, which is moistened and shaped by hand into saucer-like discoid cakes about 90 mm in diameter and 5 mm in thickness known as *agiligili*, which are then gently placed into a boiling solution of water and an alkaline wood-ash filtrate (cooking soda) and cooked into cutlets somewhat comparable in taste to sesame as the main ingredient in a rich stew, served along with cooked greens and a millet-based boiled bread (Epila-Otara, 2013). Although as the dish is occasionally enriched by the inclusion of the high-status smoked beef known as *olel*, in the author’s experience (and to his taste) this combination is overdone, since the *amola* cakes are themselves very distinctly filling for a plant-based food, evoking a pronounced satiety response.

Hyptis is situated within the foodways of the northern Uganda which derive from the Sudanic landrace crops based on foxtail millet *Eleusine coracana* (L.) Gaertn. and sorghum varieties, with a wide range of pulses including short- and long-season pigeon pea, cowpea and mung bean, as well as Bambara groundnut. Though the region has long been considered socio-economically underserved in the national context, Jameson (1958) reported the northern Ugandan diet to be highest in vegetable protein (in grams of protein per 100 calories) as compared to all other regions; Lango and Acholi were ranked first and second of the 13 ethnicities surveyed in their consumption of plant protein, and recent studies have confirmed the ongoing nutrient adequacy of dietary protein in the northern region (Marivoet & Ulimwengu, 2022).

In contemporary smallholder cultivation, hyptis is sown in polyculture with cereals and pulses, occurs spontaneously on cultivated fields, and is often conserved and tended within the residential compound, where it can be watched over as an ornamental plant and harvested for threshing when ripe with minimal loss. The northern Ugandan cultivar called *amola atar* (white hyptis) lends itself to the place name Amolatar, and is notable for its large flower heads, which have been observed to ripen simultaneously with minimal seed shattering prior to harvest. A second local variety with a black stem, called *alako*, is apparently less common (COVOL Uganda, 1997), and has not been observed by the author.

Productivity data for the crop were collected by De Schlippe (1956), who recorded hyptis yields in the Western Equatoria region of present-day South Sudan of 240 kg/ha at Yambio, 450 kg/ha at Kagelu (just south of Yei), and up to 1600 kg/ha on the alluvial soils around Maridi. An experimental trial undertaken by Teissonnier (1902) at Ditinn in the Fouta Djalon region of Guinea obtained much more modest yields of 145 kg/ha, while in a test garden closer to the Atlantic coast considered unsuitable for the species, only 3.5 kg was obtained over an area of 280 square meters, a yield equivalent to 125 kg/ha.

The first compositional studies of hyptis seed were undertaken out of interest in the crop as an oilseed. At the 1900 Paris Exposition, hyptis from the Fouta Djalon region of Guinea was featured in an exhibit of plant foods displayed in the French Guinea pavilion, leading to the first lipid analyses undertaken at Marseille (Milliau, 1901). The results, and those of later analyses of hyptis from Nigeria and Sudan, confirmed the potential of the crop as industrial feedstock based on its high content of ‘drying oils’ then in high demand – particularly linolenic fatty acid, comprising about 60% of the seed lipids, and linoleic acid at about 23% (Vuillet, 1912; The Editor, 1941). In subsequent studies, the iodine value of the oil, indicative of its drying properties, has been found to exceed 200 gI/100g – higher than the linseed oil standard, and on a par with the related species *Perilla frutescens* (L.) Britton Lamiacieae (Barker et al., 1950; Grindley 1950). From a nutritional standpoint, α-linolenic acid is a polyunsaturated fatty acid essential to the human diet due to its role in cell membrane tissue function (Burdge & Calder, 2006).

The protein content of hyptis seed was first determined in proximate compositional analysis by Grindley (1950), subsequently replicated by Leung et al. for inclusion into the FAO Food Composition Tables for Africa (1968). Compositional studies of the Bari *kino* variety cultivated in present-day South Sudan were subsequently made by de Schlippe (1956) and by Banja et al. (2016) and Abe (2019); the results of these earlier studies, indicated as percentages, are included for comparison in Table 1. Further, Banja et al. (2016) determined the anti-nutrient content of the raw seed comprising 332.0±12.98 mg/100g tannins, 434.8±57.71 oxalates and 390.4±19.60 mg/100g phytates, which were found to be reduced substantially by roasting.

**Table 1.**
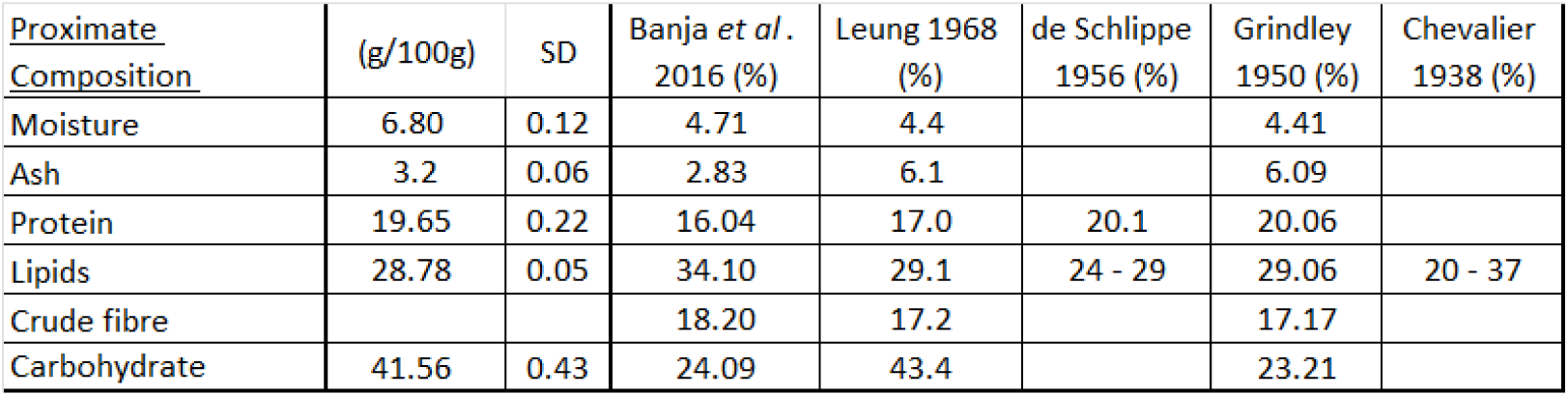
*Hyptis spicigera* proximate composition and standard deviation, with historical values.

However, no earlier study has determined the amino acids profile of hyptis, nor assessed its protein quality – notable among the relevant nutritional properties as yet undetermined for the crop, on which Banja et al. (2016) called for further research.

As a minor crop now relegated to obscurity, the historical record on hyptis is dispersed and antiquated and has not been brought together in any prior publication - leaving open the question of why, when and how the species came to be domesticated. Whereas plant domestication in Africa has been considered a response to climatic changes dated to between 10,000 and 4,500 BCE (Bellwood, 2022 p. 110), Larson et al. (2014) contend that the role of human agency and cultural factors in domestication has received little attention, and are important to understand. Hannaford (2023) argues that reviving the historical geographies of foodstuffs and food systems can provide an operative context by which to address the drivers of vulnerability to food insecurity, leveraging “usable pasts” of local knowledge, technologies and resources that provide producers with a range of options by which they may effectively respond to current conditions and future circumstances.

Relevant to its African domestication as a food crop, the taxonomic and genetic diversity within the species have not been assessed in any prior publication, although secondary data including published photographs (*e.g.* Akobundu & Agyakwa, 1998) indicate a high degree of phenotypic variation, as do the digitized herbarium specimens from international collections recently made accessible via online databases. One such database, the Global Biodiversity Information Facility, comprises 2,265 occurrences and 1,295 georeferenced records of hyptis, of which 139 entries include digitized photographs of herbarium vouchers (GBIF Secretariat, 2023). Studies of morphological diversity based on measurements obtained from such digitized herbarium specimens have confirmed the usefulness of the method (Soltis, 2017), providing broadly accurate if approximate two-dimensional values as a basis for comparison (Borges et al., 2020). Although recent work in this area has involved the use of specialized software for greater precision, a manual comparison of the specimens was made here on basis that such “quick and dirty” analysis may provide insights of practical utility, if only as a basis for further research (N. Simmonds, pers. comm. 1994).

In pursuit of these related objectives, the purpose of this study was to determine the proximate composition of hyptis seed and to assess its protein quality through amino acids analysis, while also presenting a comprehensive historical review of a precolonial food crop once widely cultivated and still highly valued by rural communities, yet largely forgotten by agricultural science.

### 2.0 Materials and Methods

In a recent paper, the author developed and tested a nutritional profiling methodology to assess the composition and protein quality of the seeds of the leguminous tree *Parkia biglobosa* sampled from three locations of Mali (Masters & Kelly, 2024). The same analytical methods were employed here to assess the composition and protein quality of a blended sample of *Hyptis spicigera* seed obtained from a rural market in the Otuke District of northern Uganda.

The proximate compositional profile of hyptis seed flour was obtained using AOAC method 935.29 for moisture analysis, 985.01 for ash, and 990.03 (combustion) with a nitrogen conversion factor of 5.3 for protein. Lipid extraction was undertaken using the FOSS Soxtech system according to manufacturer’s suggestions, following which the carbohydrates were determined by calculation (subtraction). Three replicate analyses were conducted for each test, and the results presented here as an average with standard deviation indicated for each parameter.

Amino acid profiling analyses of all samples for 19 amino acids (including cysteine, tryptophan, hydroxyproline and taurine) were performed as per the Australian Proteome Analysis Facility (APAF) standard operating procedure (SOP) AAA-001. As such, all samples underwent liquid hydrolysis for 24 hours at 110 °C in 6 M Hydrochloric acid. Under these conditions, asparagine and glutamine are hydrolyzed to aspartic acid and glutamic acid respectively, and therefore the reported amount of these acids represents the sum of those respective components. After hydrolysis, all amino acids were labelled using the Waters AccQTag Ultra chemistry following supplier’s recommendations, and then analyzed on a Waters Acquity UPLC. Four replicate analyses were conducted for each test, and the results presented as an average and a standard deviation for each parameter.

Protein quality determined using amino acid scoring, whereby the content of each essential (or conditionally essential) amino acid is compared to a reference protein to determine an Amino Acid Score (AAS), the test-specific component of the WHO/FAO Protein Digestibility-Corrected Amino Acid Score (PDCAAS), which is determined by the digestible value of the first limiting essential amino acid (Schaafsma, 2005).

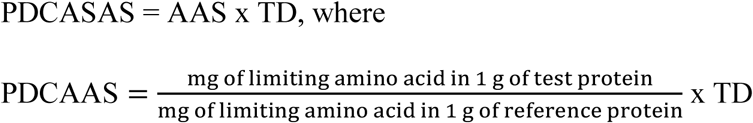

where TD is true fecal digestibility of the test protein, as measured in a rat assay.

Since actual TD values were not available, the PDCAAS of hyptis seed flour was instead tested using approximative protein digestibility (PD) values estimated at 85%, this value based on comparable figures determined for sesame (Embaby, 2010) and the edible seeds of two related species of Lamiaceae, *Salvia hispanica* L. (Beltrán-Orozco et al., 2020) and *Perilla frutescens* L. (Longvah & Deosthale, 1991), of which the latter source determined true fecal digestibility by rat assay.

Following the compositional analyses, an exhaustive review was undertaken of the earlier literature and data on hyptis in cultivation, its production and management, and its proximate composition, the results of which have been synthesized here in form of a contextual literature review.

Finally, a set of digitized herbarium specimens identified as *Hyptis spicigera* syn. *Cantinoa americana* was assembled in order to evaluate patterns of morphological variation in the shape and dimensions of the major inflorescence. Drawing from the shape typology used for leaves by Arbonnier (2004), the flower shape of each specimen was classified as linear, oblong, elliptic or ovate, orbicular, obovate lanceolate or oblanceolate. For each specimen, the length of the longest flower and its average diameter were estimated, and these dimensions were multiplied to obtain a proxy measure of flower size. Variation within these parameters was assessed and compared with respect to Neotropical, African and Asian origins.

### 3.0 Results

### 3.1 Compositional analysis

Table 1 provides the proximate composition of hyptis seed according to the standard parameters, with the historical values obtained during previous studies of different origins provided for purposes of comparison, with which the current figures are in broad alignment.

Proximate composition of the seed is 28.8% lipids and 19.7% protein, lower in lipids but comparable in protein content to some modern sesame cultivars of *Sesamum indicum* L. which are unrelated botanically (Uzun et al, 2008), while the lipid to protein ratio of 3:2 is similar to that of the wild sesame relative *Sesamum sesamoides* (Endl.) Byng & Christenh. syn. *Ceratotheca sesamoides* Endl. (Fasakin, 2004).

Table 2 presents the amino acid profile of hyptis, with standard deviation (SD) indicated for each component amino acid, including subtotal figures for essential amino acids in mg/g.

**Table 2.** *Hyptis spicigera* amino acids profile.

| Amino Acid | mean<br>(mg/g) | SD |
| --- | --- | --- |
| Alanine | 9.2 | 0.17 |
| Aspartic acid | 16.5 | 0.29 |
| Glutamic acid | 34.8 | 0.62 |
| Hydroxyproline | 1.0 | 0.05 |
| Serine | 10.7 | 0.21 |
| Arginine | 19.6 | 0.33 |
| Cysteine | 4.2 | 0.24 |
| Glycine | 9.7 | 0.17 |
| Proline | 7.9 | 0.21 |
| Tyrosine | 6.3 | 0.08 |
| Histidine | 5.4 | 0.10 |
| Isoleucine | 7.8 | 0.17 |
| Leucine | 14.0 | 0.25 |
| Lysine | 9.0 | 0.18 |
| Methionine | 2.9 | 0.17 |
| Phenylalanine | 10.3 | 0.17 |
| Threonine | 7.3 | 0.13 |
| Tryptophan | 2.6 | 0.13 |
| Valine | 9.7 | 0.17 |
| Total Amino Acids (n=19) | 188.7 |  |

Figure 2 provides a visual representation of the amino acids profile derived from Table 2, with indication of the relative proportion of essential amino acids, with SD indicated by the error bars.

**Figure 1.**
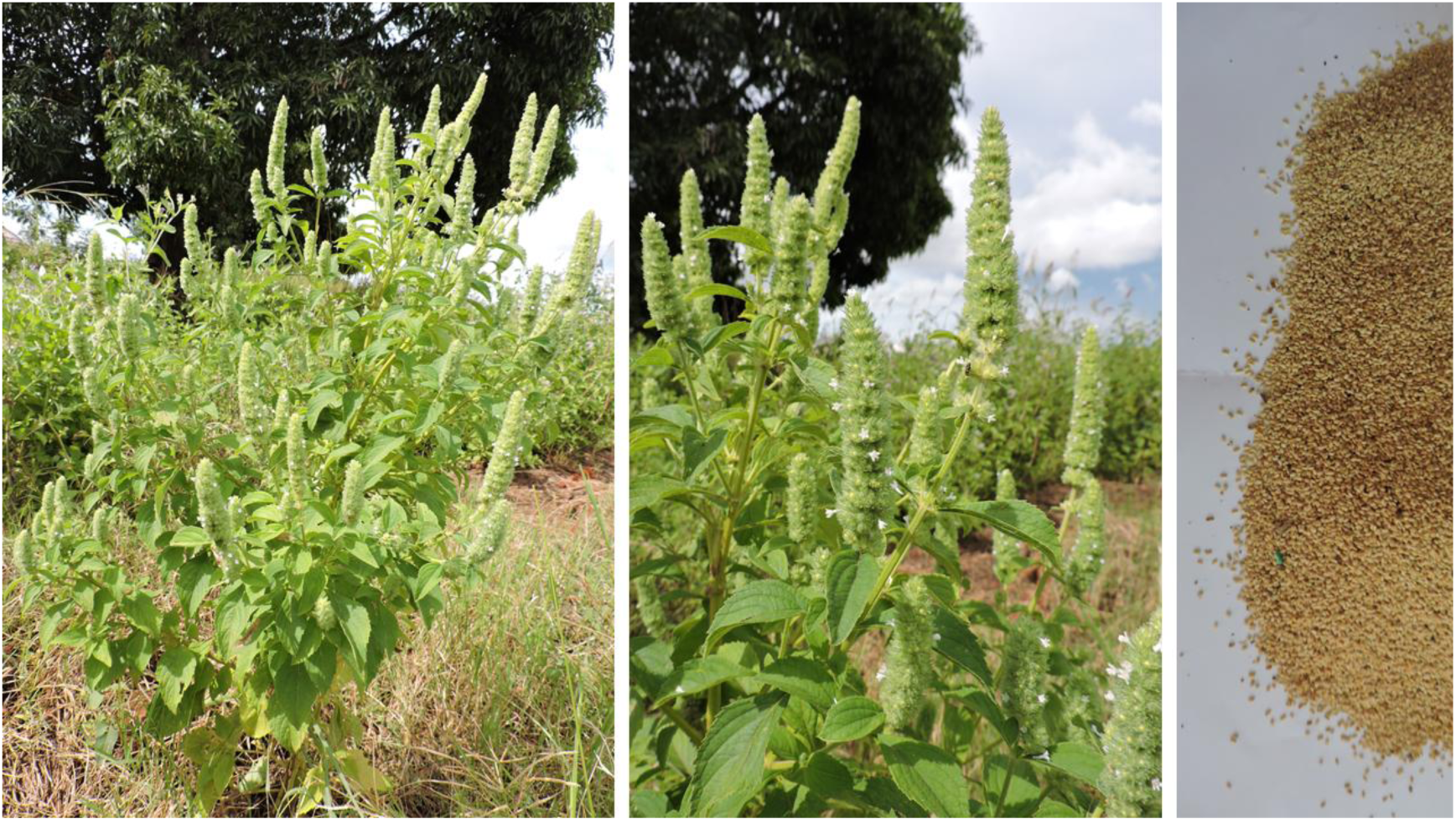
*Hyptis spicigera* (*amola atar*) a. in cultivation, b. inflorescence detail, and c. dried seed

**Figure 2.**
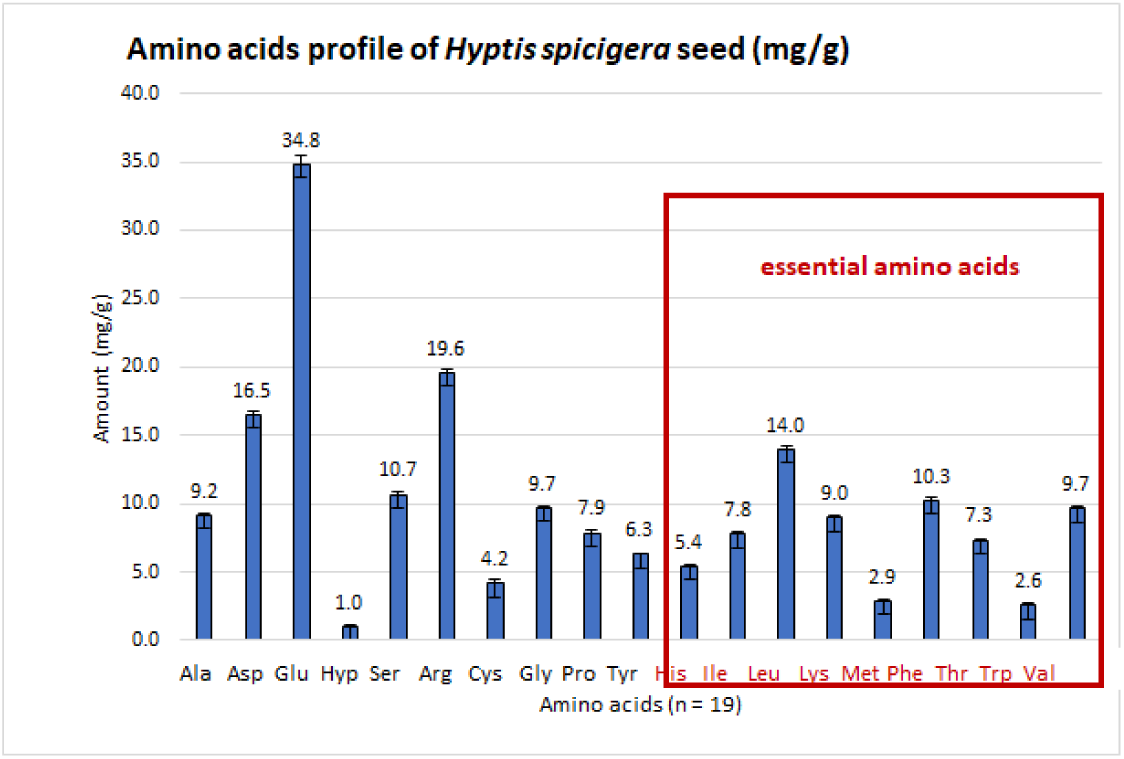
*Hyptis spicigera* amino acids profile

The amino acids profile of hyptis compares favorably to that of its early northern Ugandan co-domesticate *Hibiscus sabdariffa* L, for which the limiting amino acids have been identified to be valine, isoleucine, and tryptophan (El-Adawy & Khalil, 1994).

### 3.2 Protein quality

Table 3 presents the findings of the PDCAAS test. The limiting essential amino acid was found to be lysine at 47.7 mg/g of protein (±0.179), although the deficiency is not statistically significant, comprising 99% of the ideal Amino Acid Score (AAS) of 48 mg/g.

**Table 3.** *Hyptis spicigera* protein quality evaluation by PDCAAS.

| EAA | g of EAA in<br>100g of<br>tested<br>food | mg of EAA<br>in 1g of<br>tested<br>protein | mg of EAA<br>in 1g of<br>ideal<br>protein | AAS | PD |
| --- | --- | --- | --- | --- | --- |
| Histidine | 0.5 | 28.7 | 16 | 179 | 85 |
| Isoleucine | 0.8 | 41.2 | 30 | 137 | 85 |
| Leucine | 1.4 | 74.0 | 61 | 121 | 85 |
| Lysine | 0.9 | 47.7 | 48 | 99 | 85 |
| Methionine + Cysteine | 0.7 | 37.4 | 23 | 163 | 85 |
| Phenylalanine + Tyrosine | 1.7 | 87.7 | 41 | 214 | 85 |
| Threonine | 0.7 | 38.9 | 25 | 155 | 85 |
| Tryptophan | 0.3 | 13.7 | 6.6 | 208 | 85 |
| Valine | 1.0 | 51.3 | 40 | 128 | 85 |
| total: | 7.933 g | 319 mg | 291 mg | - | - |

The PDCAAS value of 0.84 indicates that the protein quality of hyptis seed is relatively high - comparable to many pulses and pseudocereals, and superior to both pea and quinoa (Nosworthy & House, 2017; Sá & House, 2024).

### 3.3 Contextual review of hyptis in historical observations

The historical record identifies hyptis as an important food crop in cultivation across 7,000 km the Sudanian agroforestry parkland, with nodes in the Fouta Djalon region of Guinea (the point most proximate to a transatlantic dispersal), and inland zones including Nigeria, the Adamawa Plateau of Cameroon, and the central African watershed drained by the Congo and Nile rivers, comprising present-day Democratic Republic of the Congo, Central African Republic, South Sudan and Uganda (Figure 3).

**Figure 3.**
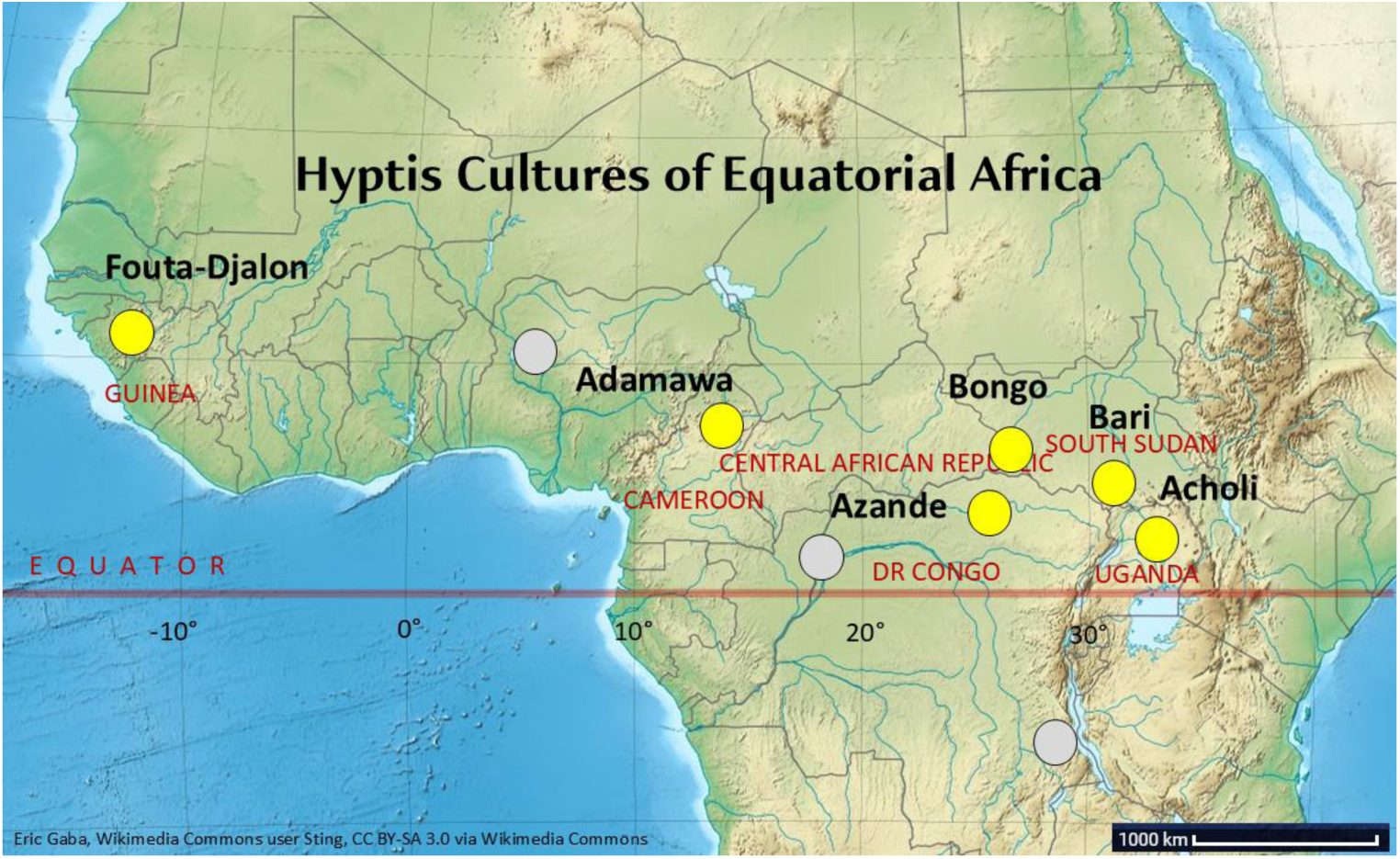
Hyptis cultures of equatorial Africa

Although not reported as a food in its Brazilian center of origin, emerged as a major crop within a few hundred years of its earliest potential arrival in Africa, having been taken up in cultivation for its protein-rich oilseed by multiple societies extending far inland from any potential port of arrival. The African domestication of hyptis occurred during historical times, and yet the timeline of its selection and diffusion as a crop is unknown - evidently predating the earliest of recorded observations, and unaddressed in the broader agricultural histories of the region (Baker, 1962; Davies, 1968; Harris, 1976; Harlan, 1982).

The species was first described by Aublet (1775) in French Guyana as *Cantinoa americana*, and in Brazil by Bentham (1832) and Gardner, c. 1846 according to Fagg *et al*. (2015).

Confirming the inclusion of *H. spicigera* (as *Pycnanthemum elongatum* Blanco) in the *Flora de filipinas* (Blanco, 1837), Merrill (1904) speculated that *H. spicigera* reached the Philippines accidentally and among several other hyptids, presumably in the form of dunnage protecting oceangoing cargoes. As of the 1830s, the species was already “common and widely distributed” throughout the Philippine islands as a weed of peri-urban waste places, while botanical observations of *H. capitata* Jacq. can be dated back to the late 1500s (Merrill, 1904). Epling (1936) described *H. spicigera* as “widely distributed throughout the East Indies, where it is more often collected than in the Americas, and in Africa where it has been found in Senegal, Sierra Leone, Nigeria, the Sudan, the Congo, Nyasa and Madagascar.” *H. spicigera* was identified as present in Guam by Epling (1936) and by Stone (1970). Dube (2017) provides, albeit without attribution, a lengthy list of local names for the species in Senegal, Mali, Burkina Faso, Côte d’Ivoire, Ghana, Nigeria, Cameroon, Indonesia and Sulawesi, and the Philippines.

In Africa, the literature identifies hyptis as a major cultivated crop at the time of the earliest recorded observations by European geographers in several sub-regions, considered as a precursor to sesame (Seignobos, 1982), with which it is botanically unrelated, yet categorically associated in the vernacular nomenclature. The prolific French tropical botanist Auguste Chevalier (1924, 1931, 1938, 1948, 1949) saw hyptis as a pantropical species proliferating in the Brazilian savanna, an American import of “several centuries” and an early famine food in Africa, naturalized in bush and fallow in Guinea, Mali and the Sudan savannahs, as well as on Madagascar and in Vietnam, “until recently” cultivated in present day Mali, Guinea, Central African Republic and the northeast corner of the (then) Belgian Congo, prior to 1910 when the crop reported (perhaps prematurely) as having been replaced by groundnut ‘or even sesame’ and its cultivation discontinued (Chevalier, 1947).

At the western extremity of its African range and as mentioned above, hyptis was well-documented at the turn of the 19^th^ century as important crop in the Fouta Djalon region of Guinea (Milliau, 1901; Vuillet, 1912); some decades later, it was found to be cultivated in rice paddy and swamps after the rice harvest in Senegal, and tended in association with upland rice in Sierra Leone (Dalziel, 1937 as cited in Burkhill, 1995). Kerharo & Bouquet (1950) found hyptis commonly cultivated like sesame as an oilseed across all of present-day Burkina Faso, and the crop was still in evidence 60 years later (Gallagher, 2010).

In Nigeria, Holland (1915) found hyptis cultivated at Nupe and Kontagora in present-day Niger State. Hyptis was an important crop of profound cultural resonance on the Adamawa Plateau of Cameroon, where Seignobos (1982) situated the species in a line of sesame analogues cultivated by the Dourou and Dowayo peoples (called ‘our ancient sesame’ by the Dourou), replacing *Polygala butyracea* Heck as the main oilseed prior to introduction of sesame under the Kwararafa confederacy (c. 1600s-1700s). Garine (2005) validated the findings of Seignobos, while pointedly expressing surprise that a crop considered as on its way out of circulation in the 1970s was still in common cultivation as a crop grown by women farmers in the Duupa mountain villages of northern Cameroon.

In present-day Democratic Republic of the Congo, De Wildeman (1903) lists several areas of hyptis cultivation including M’Toa, the site of present-day Kalemie on Lake Tanganyika.

Leihner (1983) described hyptis as cultivated in association with sorghum and finger-millet and a recognized companion to cassava in the groundnut/cassava/sesame system of the former Haut Zaïre (now Orientale province) bordering Uganda and the Western Equatoria region of present-day South Sudan.

In the areas of the present-day South Sudan bordering the West Nile region of northern Uganda, the German botanist and ethnologist Georg Schweinfurth (1873) ranked hyptis after sesame and before groundnut in order of production among the chief agricultural crops of the Bongo. During the subsequent Emin Pasha expedition, Mounteney-Jephson (1891) noted that whole granaries were used for storing hyptis seed, cultivated by the Bari as an oilseed and also eaten in the form of a porridge (pp 132 & 136). A few decades later, Broun & Massey (1929) confirmed its cultivation by the Bongo in the Fung and Bahr el Ghazal provinces, and described its role in Zande cuisine as both ‘an adjunct to stews and gravies’ and as an edible oil. Evans-Pritchard described the prominence of hyptis in mixed cultivation finger millet, sorghum, maize, cowpea and Bambara groundnuts among the Bongo (1929) and the Zande (1930) of Bahr el Ghazal and Western Equatoria. Ferguson (1948) considered hyptis a “characteristic crop of [the Yei] area… and used like sesame.” Bacon (1948) reported of Sudan that “*Hyptis spicigera* Lam. Bongo ‘kindi’, is found wild in the Fung and in Equatoria Province where it is increasingly cultivated, often mixed with dura [sorghum] and eleusine [finger millet].” Assessing the prospects for soya, then in its fifth decade of cultivation since its introduction to that region, Bacon (1948) opined that as an oilseed it “will have to do well to compare with *Sesamum* [sesame], *Arachis* [groundnut] and *Hyptis*”.

As senior research officer at the Yambio research station of the Western Equatoria region from 1948-1951, Pierre de Schlippe undertook with Azande farmers “the most detailed agricultural survey in tropical Africa yet published” according to Miracle (1967). The published results of his study provide very detailed observations of hyptis as a companion crop to finger millet in the *öti-moru* rotation with sesame and sorghum, and in association with the other Brazilian natives maize, sweet potato, cassava, groundnut alongside indigenous cucurbits and a range of pulses including cowpea, mung bean, pigeon pea and Bambara groundnut (de Schlippe,1956). According to de Schlippe, the quantity of hyptis grown was sufficient that its stems and chaff were spread across fields to be burned as a means of cleaning land for cultivation during the dry month of January. Further east, the crop is likewise known to the cross-border Bari and the Madi of West Nile region of Uganda (Katende et al. 1999, Tallantire & Goode 1975), but it is not reported from the Eastern Equatoria region of South Sudan (Grosskinsky & Gullick, 2000

The former importance of hyptis across the equatorial region during precolonial times is reflected in the abundance of its vernacular nomenclature, which categorically associates the crop with sesame, or ‘beniseed’ as it was commonly known in English at the time of colonial observations. Burkhill (1997) identifies the name *beni* as derived from a Manding root, noting that Dalziel cites the name for several species of *Sesamum* as ‘*béné*’ - from which seems to derive the name *bénéfin* for hyptis among the Mandinka of Mali (written as *bénéfin*g by Diarra et al., 2016), the Manding-Mandinka of Gambia and the Manding-Bambara of Senegal (Burkhill, 1995), recorded *téné-fi* or *bénéfin*g by the Susu of Guinea (Milliau 1901, Vuillet, 1912). In all of these languages, the term is translated as ‘black beniseed’ or black sesame. In a further, seemingly pejorative appellation, the qualifier -*dion* for slave is added by the Mandinka of Mali (-*do* by the Manding-Bambara of Senegal) as ‘black sesame of the slave’ the corresponding epithet among the Hausa of Nigeria being ‘sesame of the crocodile’ (*riidin-kadaa*) according to Burkhill (1995). The crop is also known as ‘hard sesame’ both in Sudanese Arabic as *simsim gowi* (Lado, 1989), and in the Azande language as *andaka* (Evans-Pritchard, 1960).

In northern Uganda, hyptis is *amola* to the Lango (*lamola* to the Acholi), and is sometimes called *nino* in those languages, while the related name for sesame,‘*nino muno*’ (*nino* of the white man) suggests that hyptis was the original *nino* – as Grant (1864) called it (spelled as “neeno”) prior to the adoption of sesame, with which it is categorically associated (Bedigian, 2018). Nino is likewise the name for hyptis among the Dinka who inhabit the ancestral Lwo homelands of the upper Nile (Ogot 1967), while the South Sudan Lwo call it *nina* (Banja et al., 2016).

However, it should be noted that although comparable to sesame in the composition and preparation of its seed, the strongly aromatic spike inflorescences of the hyptis plant sharply distinguish it both taxonomically and sensorially from the pendulous flowers and seed-pods of cultivated sesame. A parallel nomenclature is based on the distinctly fragrant property of the hyptis plant, which situates it squarely within the mint family Lamiaceae. For its repellent properties, the plant is called ‘mosquito bane’ (*soso guena*) by the Mandinka of Mali and Senegal, ‘ocimum [basil] of the marsh’ (*daidooyaa*) by the Hausa of Nigeria, and ‘medicine of the dead’ (*ka fulungay*) by the Dioila of Senegal (Burkhill, 1995). The biological activity of the hyptis volatile oils of have been studied using material from Burkina Faso (Lambert et al.,1985; Sanon et al., 2006; Conti et al., 2011), Mali (Sidibé et al. 2001), Benin (Boeke et al. 2004), Nigeria (Onayade et al. 1990, Ladan et al. 2010), Cameroon (Noudjou et al., 2007) and western Kenya (Othira et al., 2009), mostly identifying monoterpene and sesquiterpene compounds with its credited protection of stored grains from bruchid infestation (Onayade et al., 1990; Kini et al., 1993). Burkhill (1995) records many such non-food applications of the plant from across Africa, including the use of its flowers (fresh or burnt) to repel mosquitoes.

Medicinal uses of the plant have also documented (notably by Burkhill, 1995, drawing from earlier sources), including the use of a paste made from its leaves and flowers to treat headache in Congo (Staner & Boutique 1937) and in Ghana (Abbiw, 1990), while a tea (*tisane*) prepared from the whole plant was used to treat fever in present-day Côte d’Ivoire and Burkina Faso (Kerharo & Bouquet, 1950). Beyond these tangible applications, cultural or ritual uses of hyptis have also been reported. During an outbreak of cerebro-spinal meningitis, Evans-Pritchard (1936) reported that Zandeland was “laid under a general interdict” against eating manioc leaves or hyptis, as conveyed in a dream to a ghost diviner by order of the Supreme being, and subsequently confirmed by the poison oracle charged with investigating whether a ceremony would cause the epidemic to cease. While no similar beliefs have been recorded in Uganda, Bouquet and Debray (1974) report that a *Hyptis* species was used to ‘preserve spells and chase away spirits’ in Côte d’Ivoire.

As a recently domesticated weed, hyptis is a hardy and robust element within mixed farming systems, quick to regenerate where it was once planted, and to push up spontaneously where not. After several decades of observation, Chevalier (1947) reckoned that the species “deserves to be remembered for the speed with which it repopulates bare soil,” recommending its incorporation into fallow of 2-3 years as a mulch, while noting that it provides fuel for bush-fire (1949) - a property useful in controlled burns, as later reported by de Schlippe (1956). More recently hyptis has been screened for its potential for use in the phytoremediation of polluted Nigerian soils contaminated with hydrocarbons (Ogbo et al, 2009; Adesipo et al, 2020).

### 3.4 Morphological diversity of digitized herbarium specimens

Of 143 digitized herbarium specimens from 24 countries, 138 bore visible inflorescences for which shape could be classified and measurements obtained (for the results summarized here, please refer to the Supplementary Data file). Although the quality, condition and state of maturity of any single specimen was variable, making most individual comparisons difficult, the data taken together suggest a high degree of morphological variation characteristic of a wild and weedy species – and variation is particularly notable within the national and regional origins sampled at higher resolutions (*e.g.* Brazil, Benin, Democratic Republic of Congo, Indonesia and Philippines). While not all the vouchers examined bore legible notation, 14 African specimens were noted to have been obtained in a cultivated field fallow (including one Togolese specimen which was obtained from a cultivated field of maize and rice), while others were obtained from disturbed habitats including road verges, riverbanks and regularly flooded areas. However, only one specimen, obtained in the Labwor Hills of northern Uganda in 1963 and labelled as East Africa Herbarium voucher no. 1448 was unequivocally identified as a cultivated plant obtained from a farmed field, and may thus be considered a holotype of the cultivated variety.

The lanceolate flower shape was found in 65 of the 138 flower-bearing specimens from all origins (47%), as compared to 22% of neotropical specimens, 45% of Asian specimens, and 60% of African specimens, and its correlation with size may indicate that the parameter of shape has no greater significance. A mean 2-dimensional flower area of 4.82 cm was determined for the neotropical origins as a baseline (24 specimens), compared to which the 31 Asian specimens averaged 62% (at 3.13 cm), whereas the mean of the 75 African specimens was estimated at 6.53 cm or 128% the size of the neotropical baseline. While the 14 specimens collected in association with cultivation are larger still (at 161%), the sole Ugandan specimen bears a far larger inflorescence than any other, estimated at 620% of the baseline. At 12.5 by 2.5 cm, the lanceolate inflorescence of the cultivated Ugandan specimen is very significantly larger than any other, suggesting that as a population, the available digitized specimens are largely unrepresentative of the cultivated form of the species, with the Ugandan specimen a sole example. However, the next largest inflorescence is a lanceolate Brazilian specimen (New York Botanical Garden voucher no. 01871606 (GBIF Secretariat, 2023); although the specimen is rather sparse on the whole, it bears an inflorescence of 11 by 1.75 cm. The juxtaposition between the two specimens might suggest that the species could feasibly have been selected as a food crop at any stage of its pantropical journey, and yet this seems to have occurred only in Africa.

## 4.0 Discussion

The protein quality results seem to justify the precolonial domestication of hyptis as an oilseed and protein crop in equatorial Africa from a nutritional standpoint.

The historical valorization of hyptis as a cultivated African food crop belies its botanical identity as an American species of a humble genus of pantropical weeds, which somehow became locally valued to the extent that it came to be considered as an ancient landrace food crop - a “culturally enriching invasive species” in the felicitous phrase of Pfeiffer and Voeks (2008). The presumably rapid trajectory of the neotropical exotic species across equatorial Africa implies an active and endogenous process of precolonial agricultural innovation on the part of farmers and communities (Beinart & Middleton, 2004; Gallagher, 2016; Hannaford, 2023), with diffusion along convergent pathways of regional migration and exchange.

Although the sequence of events leading to its domestication are unknown, hyptis has long been noted among the numerous plant species which were somehow dispersed from their New World origins to sub-Saharan Africa (Baker, 1973; Roussel & Juhé-Beaulaton, 1992; Gallagher, 2016; Thibon, 2019), with ocean currents considered as a possible mechanism of transfer (Thorne, 1973; Renner, 2004; Takayama, 2008). Based on Thorne (1973), Renner (2004) proposed transatlantic dispersal via water, wind or birds - although her characterization of *H. spicigera* as “a tidal saltmarsh halophyte’ seems in error. Vuillet (1912) observes that hyptis is not cultivated within the littoral zone as it does not grow well there, and so human agency may appear to be more likely than more natural means as a vector of its transatlantic crossing.

Chevalier (1931) listed *H. spicigera* among four other hyptoids [as] “anthropophilic species (weeds) common to South America and tropical Africa” which he identified as “having been transported from one continent to another by man”. The possibility that hyptis was taken to Africa by former slaves returning to the continent is raised by comments in Ralston (1970), and Gallagher (2016), but this idea is unsupported by any known evidence, and moreover the timeline of such return would presumably have been very late with respect to the chronology of its subsequent diffusion. An earlier pantropical distribution was surmised by Merrill (1904, 1918), who speculated that *Hyptis* species had reached the Philippine islands early on in the form of cargo dunnage. Given the significant risk of seed contamination by invasive weeds in modern cultivation systems (Gervilla et al., 2019), it seems possible that hyptis could have crossed the Atlantic incidentally admixed with maize seed during the time of its introduction to Africa, perhaps among the flour maize varieties thought to have been planted along the western African littoral by the Portuguese (Portères, 1955; Willett,1962). It is known that Portuguese ships plied the coastlines of Brazil and Guinea as from 1440 (Hair, 1992), so this early date opens the historical window for such accidental transfer.

However it may have made its way to African soil, from that point forward hyptis appears to have spread rapidly and by human agency, since by 1862 it had covered 7,000 km over 40 degrees of latitude to become a major food crop cultivated by the Acholi Lwo of northern Uganda (Grant, 1864; Stapf, 1906), and in present-day South Sudan, where it was observed in cultivation by Schweinfurth shortly thereafter (1873). It is notable that these early observations were made prior to the introduction of maize, which is thought to have reached Uganda from Egypt (Portères, 1955; Miracle, 1963), remaining a minor crop in Uganda for much of the 20^th^ century (Thomas, 1970); although maize had reached Egypt as from the Ottoman conquest of 1517 (Portères, 1955), Grant (1864) pronounced it absent from Uganda as of 1862. Hyptis therefore predated maize in the sub-region, and thus the two neotropical crops arrived independently and not in association. Given that Bedigian (2018) established that hyptis (as *amola*) was not among the crops brought to the Nyanza region of Kenya in the mid-18^th^ century migrations of the Uganda Lwo, this would imply a later arrival of hyptis to Uganda, presumably from the northwest, and barely a century before Grant’s observations.

Evans-Pritchard (1960) provides anecdotal indication that hyptis was believed to have been adopted by the Zande of present-day South Sudan and Democratic Republic of Congo from the neighboring Amiangba people of the Bas-Uélé province, speculating that groundnuts and maize (along with cassava, all of Brazilian origin) reached the Zande “from the south-west”. Whitehead (1962) likewise found hyptis among the neighboring Bari of Western Equatoria region of South Sudan, in association with a maize variety he identified as being traceable to Nigeria. These observations suggest a west-to-east diffusion of hyptis across the equatorial region.

From the available botanical evidence limited to the 138 herbarium specimens, reviewed here, the high degree of morphological variation evident across the collection seems to presents the species as a weed – a visual indicator of which might include a predominantly vegetative and structural appearance, as compared to the pronounced floral predominance of the cultivated variety.

Elsewhere, the concept of ‘weediness’ has been defined as a function of “adaptation to some niche in the human habitat” and the ecological role of given species in secondary succession by which hyptis may have entered cultivation as an “encouraged weed-crop” (Harlan & deWet, 1965), per a ‘dump heap’ origin of agriculture as proposed by Sauer (1950), whereby useful weeds were among the species selected for early domestication.

The herbarium specimens include specimens from Sierra Leone and the Kongo Centrale region of present-day Democratic Republic of Congo, Atlantic locations suggestive of a West African landing from which cultivation extended eastward across the equatorial interior. At the eastern margins if its equatorial range, the densely-packed and pronounced inflorescences of the cultivated specimen from northern Uganda in particular appear indicative of the systemic, genetic and morphological changes brought about over time by human selection for favored traits in cultivation which characterize a domesticated plant (Neumann, 2005).

Geographically, this might imply a cumulative process of selection and improvement as the species spread in cultivation across the region. Whereas the inhibited seed shattering observed in the Ugandan variety is commonly considered as an indicator of plant domestication (Fuller & Allaby, 2009), it is also a trait found in some cropping weeds which are adapted for dispersal during harvest (Maity et al., 2021), and so the relevance of that trait to domestication of the species remains to be undetermined.

While the secondary data reviewed here provides some tentative insights, the genetic, phenotypic and morphological diversity of the species across its African range remains to be investigated, with particular attention to characterization of the species through elaboration of taxonomic descriptors. Genetic studies, including sequencing for gene expression profiling, could shed light on the pathways whereby the species was domesticated in cultivation, and the precolonial cross-regional exchanges whereby the crop was adopted (Zsögön et al., 2022). Although the domestication process reduces biodiversity by definition, Krug et al. (2023) propose that future adaptive solutions may be derived from a better understanding of the agricultural past, toward which it is hoped that this study may contribute.

## 5.0 Conclusions

While the proximate compositional figures obtained broadly align with earlier published data, the amino acid profile presented here provides the first such analysis and assessment of the edible protein quality of the *Hyptis spicigera* seed, which was determined to relatively high.

In historical context, these results shed light on the nutritional value of a precolonial food crop, now largely forgotten, which should be recognized among the plant-based protein foods of the sub-Saharan cultivation systems (Ogutu et al., 2024).

Based on the observed of the traditional dish prepared from the seed, further studies should explore its organoleptic attributes of hyptis as a food, including its satiating properties (Holt et al., 1995, Forde et al., 2013), a parameter of relevance in the search for novel plant foods and plant-based protein products (Chambers et al., 2015; Javanmardi et al., 2021).

It is hoped that these results may serve to raise the profile of a largely forgotten food crop with a rich history reflecting precolonial agricultural innovation and exchange. Although hyptis presents the curious case study of a little-known American weed that became an African food crop, it is but one among a wealth of locally-adapted landrace food crops maintained in cultivation by smallholder farmers within a little-studied ‘non-center’ of crop diversity, all of which merit further attention and policy support to ensure their availability and survival into an uncertain agricultural future.

## Supporting information

Supplemental data morphology

## Data availability

The analytical data upon which this study is based will be shared with all interested researchers upon request.

## Acknowledgments

This paper is dedicated to the memory of Jaspher Okello, who undertook sampling from which this study was based and passed away unexpectedly during the background research phase of publication.

Special thanks to Perpetua Apili for her organization of field visits in Agago and Otuke districts of northern Uganda during October 2019, and to Leonora Okello, Pirina Okidi and Daniela Oyaka of Agago District; Grace Abuka of Corner Adwari, Otuke District and Ladit Bejaleri Ongura of Alula, Acan Pii (Otuke County) for hosting visits to their gardens and homesteads in which hyptis was in full flower.

The author likewise extends his sincere appreciation the women farmers who hosted and fed him so well during the earliest phases of his research him in the rural areas of Otuke (1990-1996), during which time he learned to appreciate *amola* as a special dish. Thanks also to the young researchers of the Northern Uganda Food Security Project (DERO) and the Shea Project (1997-2000), who investigated hyptis as a landrace crop of cultural significance.

The author recognizes with gratitude the contribution of the NMIT Research Trust for funding of the sampling and analysis phases of this study. Proximate compositional analysis was undertaken by the Food Processing Center at the University of Nebraska-Lincoln, and the study used the NCRIS-enabled Australian Proteome Analysis Facility (APAF) infrastructure at Macquarie University, which undertook amino acids analysis. The author is likewise grateful to these study partners for their analytical support.

